# An interpretable deep learning framework for genome-informed precision oncology

**DOI:** 10.1101/2023.07.11.548534

**Authors:** Shuangxia Ren, Gregory F Cooper, Lujia Chen, Xinghua Lu

**Affiliations:** Intelligent Systems Program, School of Computing and Information, University of Pittsburgh, Pittsburgh, Pennsylvania, United States of America; Department of Biomedical Informatics, School of Medicine, University of Pittsburgh, Pittsburgh, Pennsylvania, United States of America

## Abstract

Cancers result from aberrations in cellular signaling systems, typically resulting from driver somatic genome alterations (SGAs) in individual tumors. Precision oncology requires understanding the cellular state and selecting medications that induce vulnerability in cancer cells under such conditions. To this end, we developed a computational framework consisting of two components: 1) A representation-learning component, which learns a representation of the cellular signaling systems when perturbed by SGAs, using a biologically-motivated and interpretable deep learning model. 2) A drug-response-prediction component, which predicts the response to drugs by leveraging the information of the cellular state of the cancer cells derived by the first component. Our cell-state-oriented framework significantly enhances the accuracy of genome-informed prediction of drug responses in comparison to models that directly use SGAs as inputs. Importantly, our framework enables the prediction of response to chemotherapy agents based on SGAs, thus expanding genome-informed precision oncology beyond molecularly targeted drugs.

## Introduction

Precision medicine utilizes genomic and other advanced technologies to define diseases at a more detailed level than before, enabling tailored therapies for individuals^1,2^. This approach largely relies on understanding the impact of genomic alterations within cells and prescribing medications to counteract aberrant signals caused by these alterations. The common practice of genome-informed precision oncology is to examine the somatic genome alterations (SGAs) and match patients with targetable SGAs to corresponding targeted drugs ^1,3,4^. While of clinical value, this approach is applicable to a relatively small number of molecularly targetable drugs, patient coverage is relatively low, and prediction accuracy (positive predictive value) remains modest ^5-7^. Marquart et al ^5^ reported that as of 2018, the percentage of patients who receive genomic screening and could be matched with targeted therapies was only about 15%; the median overall response rate to all genome-informed therapies was 54%; and the percentage of all cancer patients estimated to benefit was about 7%. Thus, the current practice is insufficient to meet the needs of precision oncology for the general cancer population.

Although chemotherapies remain the backbone of general oncology, their application is largely not guided by genomic information. Recently, Liu et al ^8^ systematically studied mutation-treatment interactions based on real-world patient data and discovered that certain mutations are associated with responses to certain chemotherapy agents. Generally speaking, a “mutation-to-treatment” rule for guiding molecularly targeted or chemotherapeutic agents fails to consider that multiple SGAs in a cancer cell may influence the cellular state and, thereby, drug responses, which may contribute to the observed low accuracy ^5^ of the current genome-informed precision oncology. Thus, there is an urgent, unmet need to develop comprehensive clinical decision support systems (CDSSs) capable of utilizing genome-scale omics profiles of tumors to guide the selection of effective anticancer drugs from the entire pool of FDA-approved agents.

Developing a CDSS for guiding all anticancer drugs in pan-cancer patients using real-patient data remains challenging because it would require large-scale randomized trials testing many drugs in all cancer types, which is not feasible. To address the challenge, large-scale pre-clinical models screening anticancer-drug sensitivity have been developed by the Genomics of Drug Sensitivity in Cancer (GDSC) ^9,10^ and the Cancer Cell Line Encyclopedia ^11^. The GDSC project has examined multi-omics profiles of close to a thousand cancer cell lines and recorded their response to hundreds of drugs. This dataset fills the gaps for developing artificial intelligence (AI) models for pan-cancer and pan-drug precision oncology. GDSC studies indicate that transcriptomes of cell lines are more informative features than SGAS in predicting cell line drug sensitivity. However, in clinical practice, genomic data are more readily available, and thus effectively utilizing such information would be of high clinical value. Therefore, we set out to develop a computational framework to predict drug sensitivity based on SGA data of cell lines.

Developing a genome-based CDSS faces several challenges: 1) Drug responses are usually determined by the state of multiple signaling pathways in a cancer cell. Therefore, the genomic status of individual genes considered in isolation is insufficient to predict drug sensitivity; 2) A signaling pathway can be perturbed by SGAs affecting different member genes in the pathway that bear similar consequences on drug responses; and, 3) the SGAs perturbing a common signaling pathway tend to be mutually exclusive in individual tumors ^12,13^. As such, the signal of one SGA on a drug response may become noise when training a model learning the signal of another SGA on the same drug.

To overcome the above challenges, we developed an AI system that first transforms the SGA data of cancer cells into a representation of cellular signaling systems and then learns to predict the drug responses of the cells based on the inferred cellular states. The framework consists of two main modules: 1) A representation-learning module using the <u>Res</u>idual <u>G</u>enome Impact <u>T</u>ransformer (ResGit) model (**Fig. 1C**), which infers the cellular states based on the SGAs of a cancer cell line, and 2) a drug-response-prediction module (**Fig. 1D**), which predicts the cells’ responses to drugs based on the inferred cellular states. The combined system is referred to as the <u>ResGit</u>-based <u>D</u>rug <u>R</u>esponse Prediction (ResGitDR) model (**Fig. 1A**). We show that by more closely mimicking the cellular signaling systems, the ResGit model can learn interpretable and biologically sensible representations of the impact of SGAs on cellular signaling systems. We also show that by considering cellular states, the ResGitDR performs better in predicting drug response to both molecularly targeted and chemotherapy agents than the models that only use SGAs as inputs. Finally, we show that ResGitDR indeed takes advantage of the cellular states learned within our framework and performs state-oriented predictions. The results presented below support that the ResGitDR framework provides a new and promising direction for developing biologically motivated and interpretable systems for predicting drug responses.

**Fig. 1.**
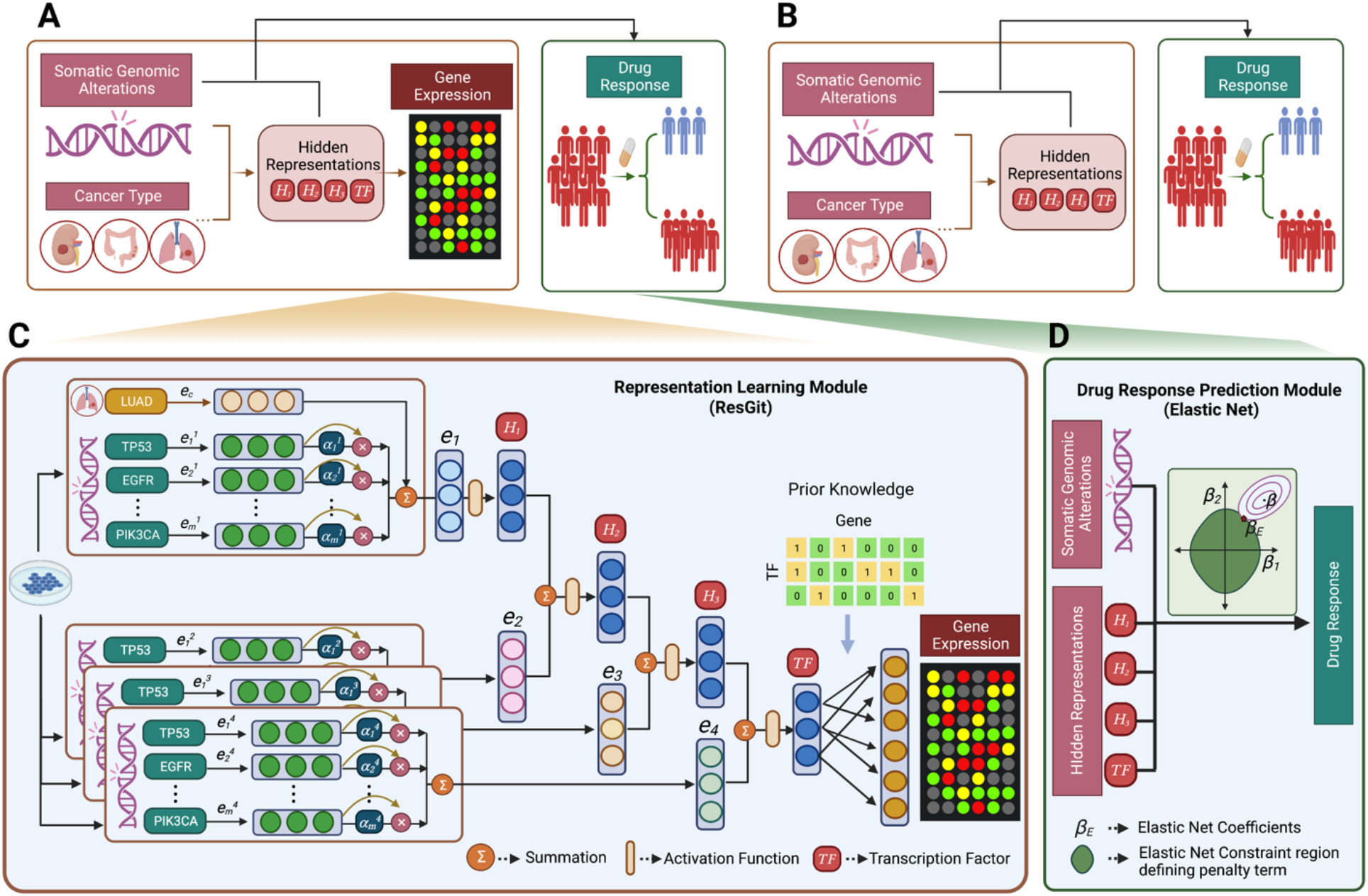
Flowchart of overall drug sensitivity prediction framework. **(A).** ResGitDR comprises two modules: the Representation Learning Module, which employs the Residual Genome Impact Transformer (ResGit) model, and the Drug Response Prediction Module, which utilizes an elastic net. In the training phase, the Representation Learning Module uses SGAs and cancer types to predict gene expression, and the Drug Response Prediction Module incorporates the hidden representations learned in the Representation Learning Module and SGAs as input to predict drug sensitivity. **(B).** In the testing phase, the trained ResGit model is used to obtain hidden representations using SGAs and cancer type as input. These hidden representations are then combined with SGAs as inputs to predict drug response. **(C).** The detailed diagram of the Representation Learning Module. **(D).** The detailed diagram of the Drug Response Prediction Module.

## Result

### Overview of the ResGitDR model

Heterogeneous responses to a drug by different cancer cells can be attributed to the heterogeneity of cellular states, which are driven by distinct causal SGAs that perturb cellular signaling systems. Thus, the capability of inferring cell states of cancer cells based on their SGAs lays a foundation for predicting drug responses. Based on the assumption that driver SGAs eventually influence gene expression, we designed ResGit (**Fig. 1C**) to model the relationships between SGAs and gene expression. It uses hierarchically organized latent variables to represent the cellular signaling system of cells and encode the impact of SGAs^14^. It then transforms the encoded information to predict gene expression.

Specifically, for each tumor, a binary vector indicating which genes are perturbed by SGA events is fed into ResGit to predict gene expression. Then four distinct embedding layers are applied to convert the binary vector into four hidden-layer-specific SGA embedding matrices, which represent the impact of SGAs in a tumor on the signal-encoding hierarchy. Each SGA embedding matrix is fed through a multi-head self-attention component to derive tumor-specific signal embedding (*e_i_*), representing the integrated impact of SGAs in a tumor on the signaling systems. The state of an internal hidden layer (*H_i_*) is a function of signal embedding (*e_i_*) and the state of the previous layer (*H_i-1_*). To incorporate the knowledge of transcription factors (TFs) on gene expression, we instantiated the final hidden layer based on prior knowledge following the example by Tao et al ^15^, such that the parameters associated with known TF-gene edges are updated during training, and the rest is set to 0. ResGit is trained with SGA and expression data of TCGA tumors and GDSC cell lines. To predict the drug response, we trained an elastic network model ^16^ for each drug. We combined the inferred state of the latent variables (reflecting cellular states) from ResGit and SGAs of cell lines as inputs and binarized drug sensitivity as the target (**Fig. 1D**). In the testing phase, as shown in **Fig. 1B**, the trained ResGit model is firstly used to obtain hidden representations by taking SGAs and cancer type as input, no gene expression data is needed during this process. Then these hidden representations are then combined with SGAs to predict drug response.

### ResGit learns to encode the impact of SGAs and transforms it into the gene expression of tumors and cancer cell lines

We collected SGA and gene expression data from 8,586 TCGA tumors and 976 cancer cell lines studied by GDSC. We trained the ResGit model using this combined dataset through a series of experiments. We evaluated model performance using the Spearman correlation coefficients between predicted and observed gene expression values of a gene as the performance metric. The distributions for the coefficients in different cancer types are shown as box plots in **Fig. 2A&B**. The mean correlation in TCGA is 0.8, while in GDSC is 0.72. The results indicate that ResGit can accurately map SGA input data to gene expression predictions. The results support that the latent variables in the model encode the impact of SGAs on the cellular signaling system and translate the information of SGAs to gene expression. Interestingly, when modeled separately, the GDSC dataset exhibited lower Spearman correlations than the TCGA dataset, which suggests that the larger sample size in the TCGA dataset made the prediction more robust, resulting in higher correlation values. From here on, we report the results of ResGit trained with pooled TCGA and GDSC data.

**Fig. 2.**
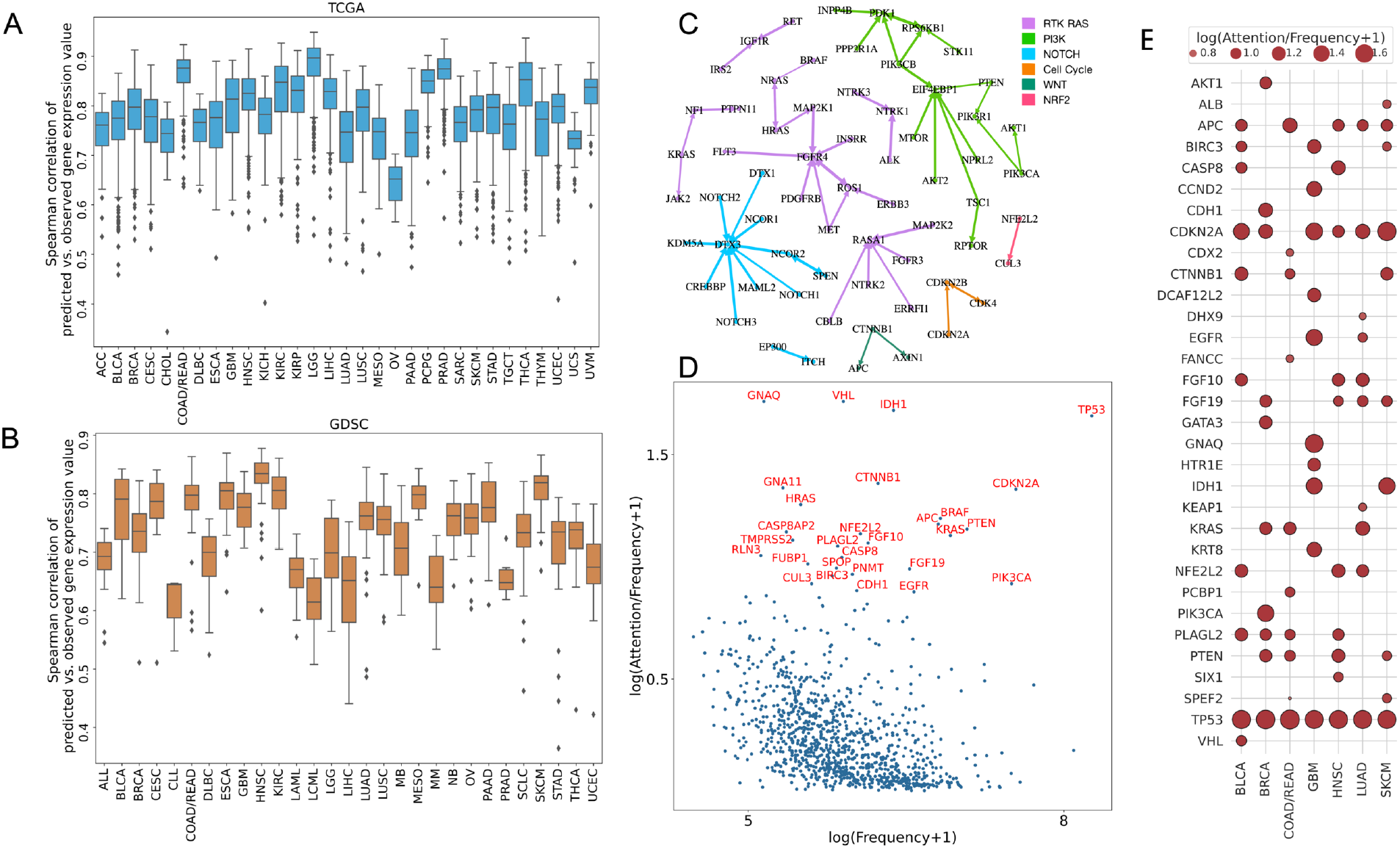
Evaluation of the performance of ResGit. The distribution of Spearman correlation coefficients between predicted and observed gene expression values **(A)** in the TCGA dataset and **(B)** in the GDSC datasets, respectively. (**C)**. The connectivity map shows the similarity of SGA embeddings among the SGAs perturbing common pathways. The weight vector connecting an SGA to hidden nodes is used as an embedding of the SGA, and similarity between a pair of SGAs is calculated with cosine similarity. If gene *A* is a neighbor of gene *B*, the arrow direction points from gene *B* to gene *A*; a double-arrowed edge indicates that two SGAs are mutually among the top 10 neighbors. The thickness of an arrow represents the degree of similarity. **(D).** The attention weights of SGAs gene in a pan-cancer analysis. Genes with high overall attention weights are shown in red font. **(E).** The attention weights of SGAs gene across different cancer types.

### ResGit captures biologically sensible representations of SGAs

In the ResGit model, an SGA is designed to be connected to every latent variable in the signaling hierarchy, and SGA embeddings represent the impact of the SGA on the system, and the model learns “optimal” connections between SGA and hidden nodes that would predict gene expression well. If two SGAs affect distinct members of a common pathway, their impact on the cellular signaling system should be similar, i.e., their embedding should be similar. We examined all pairwise similarities of SGA embeddings using cosine similarity. We identified the top 10 neighbor SGAs for each SGA and examined whether they perturb a common signaling pathway according to existing knowledge.

Sanchez-Vega *et al*.^17^ had reported SGAs perturbing ten major cancer pathways, which was used as ground truth for evaluating our results. We constructed a connectivity graph among 64 SGAs gene found in both our dataset and the reported cancer pathways gene by Sanchez-Vega *et al*., where an edge was added between a pair of SGAs if one (or both) of them was among the neighbors of the other. We colored the edges with a pseudo-color corresponding to a pathway if the connected SGAs were in a pathway (**Fig. 2C**). The learned embeddings of the members of the PI3K pathway *PIK3CA, PIK3R1, PTEN,* and *AKT1* are among the closest neighbors to each other. The graph also shows similar results for other cancer pathways. The results indicate that ResGit has learned embeddings of SGAs reflecting their similar impact on cell signaling systems, conforming to established knowledge.

### Self-attention mechanism revealed the impact of SGAs in cancers

ResGit employs self-attention mechanisms and assigns a tumor-specific attention weight to an SGA observed in a cell line to reflect its relative importance. Collective attention assigned to an SGA reflects its importance in influencing gene expression in cancers (**Fig. 2D**) or in different cancer types (**Fig. 2E**). As shown in **Fig. 2D**, ResGit assigned high attention values to well-known cancer drivers^18^, such as *TP53*, *PTEN*, *KRAS*, *BRAF*, etc. Interestingly, some genes encoding signaling proteins, such as G-proteins *GNAQ* and *GNA11,* are not well-known as “cancer drivers” but were assigned with high attention weight, despite their relatively low frequencies. The results suggest ResGit captures their impact on gene expression of cells and potential role in cancers, which is supported by recent research indicating they may play an essential role in the tumorigenesis ^19^. Our analysis also revealed the importance of SGAs in different cancer types (**Fig. 2E**). For example, the results show that SGA events in *GATA3* play a significant role in breast cancer (BRCA), as confirmed by Takaku *et al*. ^20^; SGAs in *DHX9 and KEAP1* appear to play a significant role in lung cancer (LUAD), aligning with previous studes^21,22^; alterations in *TP53* are universally involved in most cancers, as demonstrated by earlier research^23^.

### The latent representation of the cellular system is informative of drug sensitivity

The results above indicate that ResGit can encode the signals perturbed by the SGAs using the latent variables in the deep learning model. We then set out to test whether the information represented by the latent variables can be used to predict cancer cell responses to anticancer drugs.

As a baseline, we used SGAs and cancer-type labels as input to train an elastic network model (EN, **Supplementary Fig. S1A**) and an end-to-end feedforward neural network (NN, **Supplementary Fig. S1B**) model to predict cell sensitivity to each drug tested by GDSC. We evaluated the performance of each model in 10-fold cross-validation experiments. The EN and NN models for 367 drugs achieved moderate performance in terms of area under the receiver operating curve (AUROC) (**Fig. 3A**), with median AUROC at 0.595 and 0.619 for the NN and EN, respectively. We arbitrarily set the threshold that an AUROC of 0.7 indicates a potentially useful model in the clinical setting. The total number of models with AUROC above 0.7 is 7 and 32 for NN and EN, respectively. Interestingly, in this setting, the elastic network outperforms the neural network model, suggesting it is more robust in a setting with a small training sample size.

**Fig. 3.**
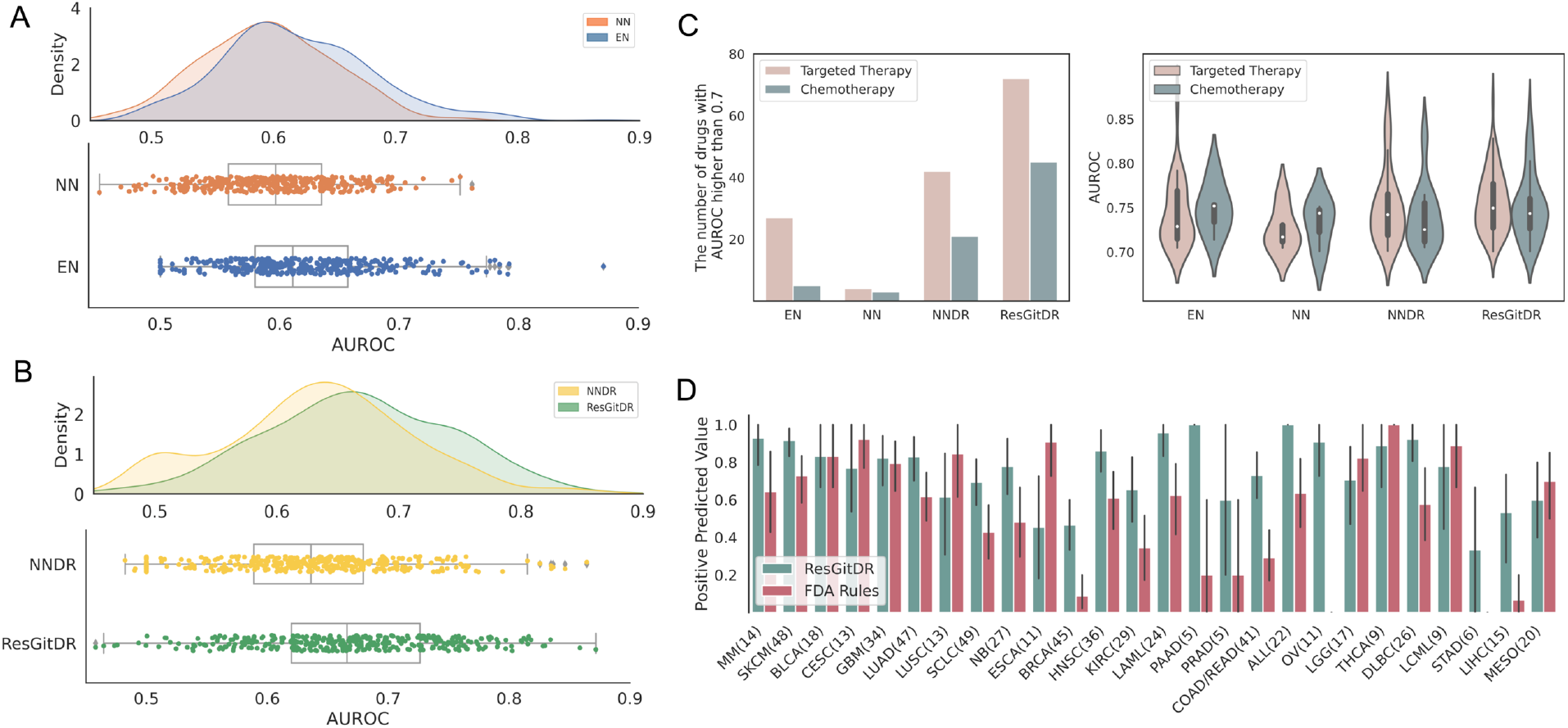
The performance comparison in drug response prediction. **(A).** The performance of two baseline models (EN and NN) which both use SGA and cancer type to predict drug sensitivity directly. **(B).** The performance of two models (NNDR and ResGitDR), which both firstly use the SGAs and cancer type to predict gene expression and obtain the hidden representations, then concatenate SGA and hidden representations to predict drug sensitivity. **(C).** The number and AUROC distribution of Targeted Therapy and Chemotherapy drugs with AUROC higher than 0.7 across EN, NN, NNDR and ResGitDR. **(D).** The Positive Predicted Value of ResGitDR and FDA rules methods with error bar representing 95% confidence interval. The numbers in parentheses indicate the corresponding cell line counts for each cancer type.

We then examined whether the latent representation learned by ResGit is informative with respect to drug sensitivity. In a 10-fold cross-validation experiment, we trained ResGit and retrieved the estimated states of latent variables (*H_1_* – *H_3_, and TF*, **Fig. 1C**) for the GDSC cell line in the training dataset. We concatenated the states of the latent variables with the original SGAs of each cell line as input features and trained an elastic network model for each drug (**Fig. 1D**). We called these models the ResGit-based Drug Response prediction model (ResGitDR). To examine the value of self-attention and other unique approaches of ResGit, we also trained a conventional neural network to model the relationship between SGAs and gene expression without direct connections from SGAs to internal latent nodes or self-attention. We extract the estimated hidden-node states to train an elastic net model, and we call this model the neural-network-based drug response prediction model (NNDR, as shown in **Supplementary Fig. S1C**). The median AUROCs of the models are 0.667 and 0.633 for ResGitDR and NNDR, respectively (**Fig. 3B**), which are significantly higher than EN and NN (ResGitDR vs. each of the rest, p < 0.01).

The numbers of models with AUROC greater than 0.7 are 117 and 63 for the ResGitDR and NNDR, respectively, and the detailed information about these drugs are listed in **Supplementary Table. S1**. Compared to the EN model, which only uses the original SGAs and cancer type as features, including the states of latent variables in ResGitDR and NNDR led to 3.7 and 2-fold increases in the number of models with AUROC greater than 0.7. We further examined models’ performances for targeted therapy and chemotherapy drugs by the four methods as an indication of what information is provided by input features and captured by the models (**Fig. 3C**). The number of ResGitDR models for targeted therapy agents with an AUROC larger than 0.7 is 72, which is 1.7-fold that of NNDR and 2.7-fold that of the EN model. Importantly, the results show that for many chemotherapy drugs, ResGitDR achieved comparable performance in terms of AUROC when compared with molecularly targeted drugs. The number of ResGitDR models for chemotherapy drugs with AUROC above 0.7 is 45, which is 2.1-fold of NNDR and 9-fold of the EN model. The results indicate that it is possible to perform genome-informed precision chemotherapy, beyond molecularly targeted drugs.

To examine the potential clinical utility of ResGitDR, we performed a simulated clinical decision experiment of assigning FDA-approved drugs to cell lines based on FDA guidelines and compared it with decisions by ResGitDR. There are 61 FDA-approved drugs (different drug_id in GDSC), 39 are for targeted drugs, and 22 are for chemotherapy agents. We applied the FDA guidelines based on cancer types and genomic biomarkers, with a preference for targeted therapy over chemotherapy. For example, the targeted therapy lapatinib is assigned to LUAD cell lines hosting SGAs in *EGFR*. If multiple drugs are eligible for a cell line, we select the one with the highest response rate among cell lines of a given cancer type, with a preference for targeted drugs over chemotherapy ones. We compared the positive predictive values (PPVs) of simulated FDA-guideline-based decisions and ResGitDR decisions.

As shown in **Fig. 3D**, in the majority of cancer types, such as MM, SKCM, LUAD, SCLC, NB, BRCA, HNSC, KIRC, LAML, PAAD, PRAD, and OV, ResGitDR predictions would make better recommendations on average. The FDA rules perform better than ResGitDR in a few cancer types, such as CESE, LUSC, ESCA, LGG, THCA, LCML, and MESO. The average PPV across all cancer types for ResGitDR and FDA rules are 0.761 and 0.549, separately. Interestingly, all OV cell lines have *BRCA1* and/or *BRCA2* mutations, and rucaparib was assigned to these cell lines per FDA rules, but these cell lines didn’t respond to this drug, leading to a PPV of zero. Similarly, cell lines in STAD were assigned with sunitinib according to the above rules and got zero positive predicted value.

To illustrate the utility of our two-component framework of first learning representation of cellular systems using gene expression as objectives and then performing cell-state-oriented drug-response prediction, we also trained a model with the same architecture as ResGit to predict drug sensitivity directly, referred to as SGA2DR model (**Fig. 4A**). The performance of SGA2DR model was worse (mean AUROC 0.602) (**Fig. 4C**) than that of ResGitDR, indicating that learning relationships between SGA and gene expression led to a better representation of cellular states that enhanced the performance of downstream drug sensitivity prediction.

**Fig. 4.**
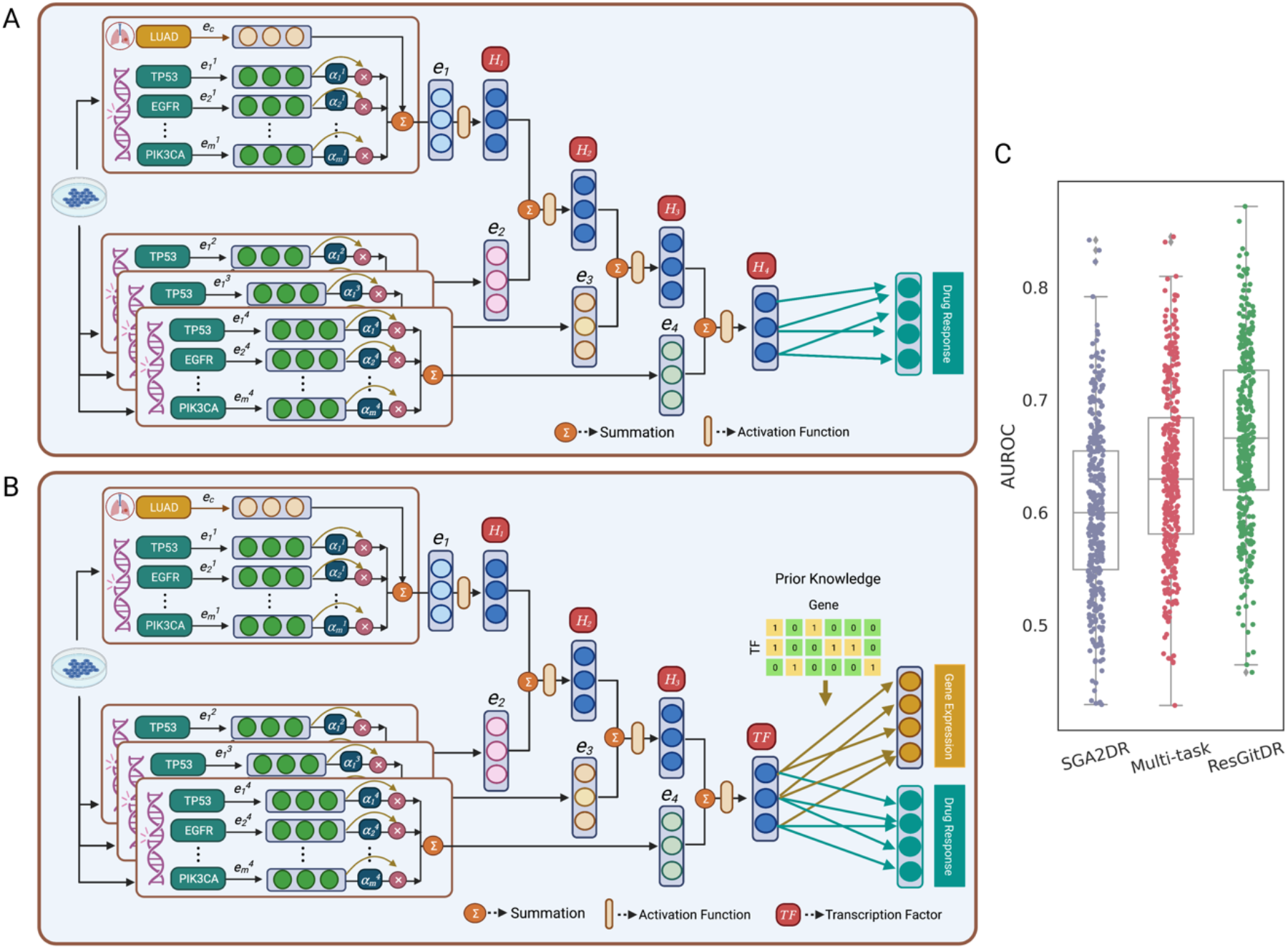
**(A).** The architecture of the SGA2DR model. It predicts drug sensitivity directly using the same architecture of ResGit by taking the cancer type and SGAs as input. **(B).** The architecture of the multi-task learning model. It aims to predict drug sensitivity and gene expression simultaneously using the same architecture of ResGit by taking the cancer type and SGAs as inputs. **(C).** The performance comparison of SGA2DR, multi-task learning, and ResGitDR models.

Further, we trained a multi-task learning model, which aimed to predict gene expression and drug response simultaneously (**Fig. 4B**). Interestingly, this model performs better (mean AUROC 0.635) than the aforementioned SGA2DR model, indicating that including gene expression as an object led to a better representation that enhanced drug response prediction. However, the multi-task model’s performance was inferior in predicting drug sensitivity compared to the two-stage approach of ResGitDR (**Fig. 4C**). This could be due to the limited size of our dataset, which consisted of only around 1000 samples. With its increased number of parameters, the multi-task model is prone to overfitting.

Finally, as a control, we shuffled SGAs and cancer-type data and re-trained a ResGitDR to predict drug sensitivity. As anticipated, the average AUROC dropped to 0.5, indicating that ResGitDR captures the “true” impact of SGAs and cancer type, which is required for predicting drug response (**Supplementary Fig. S2)**.

### ResGitDR predicts responses to molecularly targeted drugs in a cell-state-oriented fashion

Contemporary genome-informed precision oncology assigns treatment based on the genomic status of targeted signaling proteins. We evaluated the utility of genomic biomarkers for drugs targeting the PI3K/mTOR pathway, more specifically, PIK-93 and AKT inhibitor VIII, by examining whether cell lines carrying SGAs in these member genes are more sensitive (lower IC50s) than general cell lines (**Fig. 5A&B**). The results show that none of the SGAs in the pathway is informative of the sensitivity of the drugs when measured by IC50, whereas the cell lines predicted to be sensitive to the drugs by the ResGitDR models exhibit significantly lower IC50 (more sensitive). The results suggest that by considering the inferred cellular states, ResGitDR performed better in predicting molecularly targeted drugs than the conventional genomic biomarkers.

**Fig. 5.**
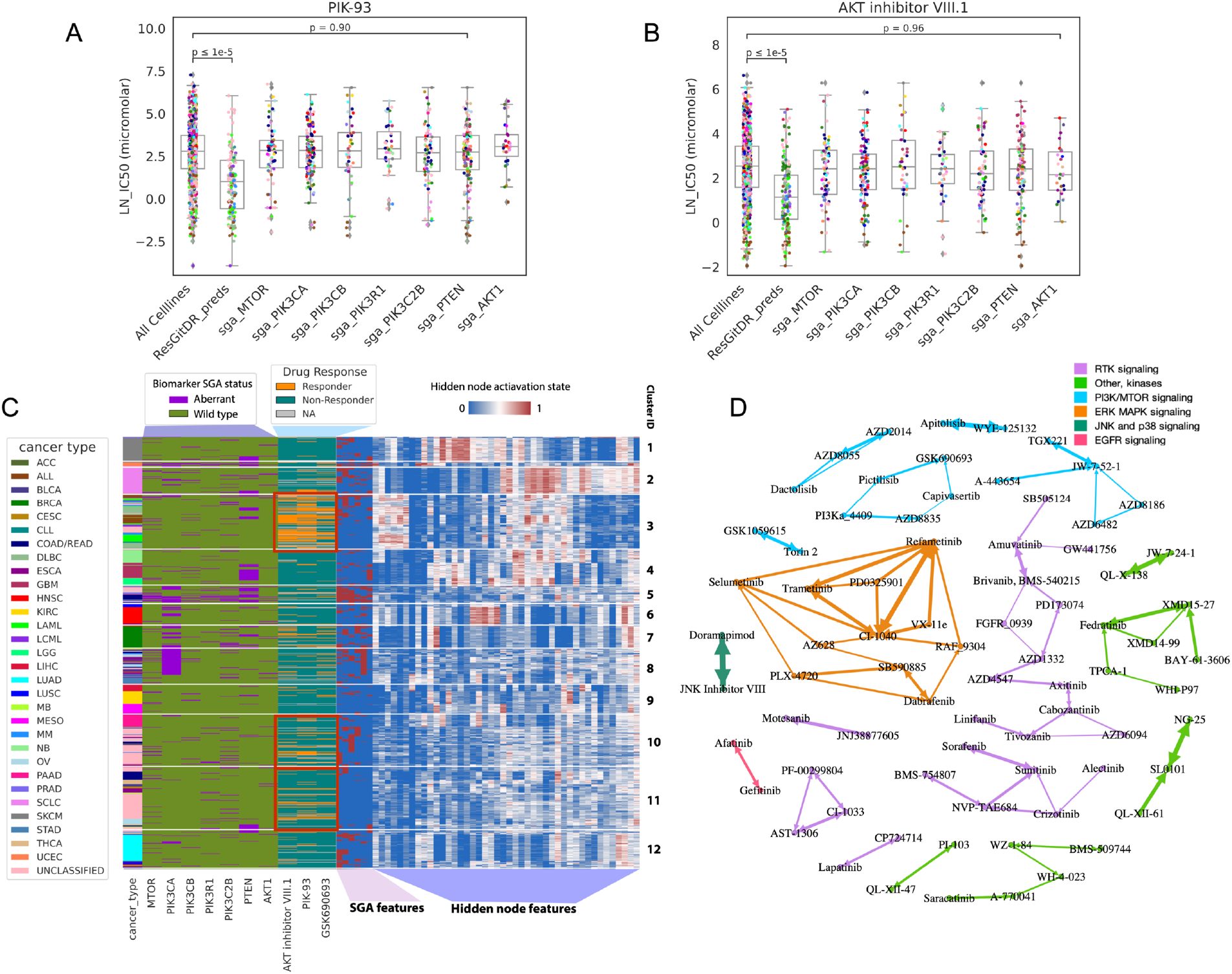
The distributions of drug sensitivity (represented as log IC50s) to **(A)** *PIK-93* and **(B)** *AKT inhibitor VIII* by cancer cell lines grouped according to the mutation status of genes involved in the PI3K pathway. The distribution of drug sensitivity by the cell lines predicted by ResGitDR to be sensitive to the drugs is also shown. **(C).** Cancer cell lines were clustered using the based on the selected top 50 predictive features from ResGitDR models for 3 anti-PI3K PI3K/MTOR drugs: *AKT inhibitor VIII*, *PIK-93*, and *GSK690693*. The features consist of hidden representations and individual SGAs. The SGAs are represented as binary values. The hidden node values are standardized within the range of 0 to 1. The binary drug responses to each of the three drugs by cell lines are shown. Three red boxes highlight the clusters with enriched responder cell lines (clusters 3, 10, and 11). The mutation status of genes in the PI3K/mTOR signaling pathway is shown to illustrate their relationship with respect to drug sensitivity. **(D).** The connectivity map shows the similarity of the embedding of drugs targeting common pathways. The top 50 important features of the ResGitDR for a drug are used as its embedding. The similarity of embeddings of two drugs is measured with cosine similarity. Molecularly targeted drugs are shown as nodes; an edge is added between a pair of drugs whose embeddings are among the top 5 highest cosine similarities of each other. If drug A is a neighbor of drug B, the arrow direction points from drug B to drug A; a double-arrowed edge indicates that a pair of drugs are mutually among the top 5 neighbors of each other. The thickness of an arrow is proportional to cosine similarity.

We then investigated whether ResGitDR utilized certain characteristic cellular states to predict responses to drugs that share similar mechanisms of action (MOA), e.g., drugs targeting the PI3K/mTOR pathway. We extracted the parameters from the models for three drugs, *ATK inhibitor VIII.1*, *PIK-93*, and *GSK690693*, and we identified a union of the top 50 features based on the absolute weights of drugs targeting on PI3K/mTOR pathway in the elastic net model, which reflect the importance of a feature, including both hidden representations and SGAs. We extracted the values of these features from GDSC cell lines and grouped them using clustering analysis (**Fig. 5C**). The cell lines’ mutation status of genes in the PI3K/mTOR signaling pathway is shown to illustrate whether they carry information with respect to drug sensitivity as biomarkers. The figure shows that inferred cell states underlie cell line clusters consisting of cells from diverse cancer types, and certain clusters (e.g., clusters 3, 10, and 11) are enriched with responders to the three drugs, supporting the notion that cell states influence the response to drugs. The AUROCs for the three models are 0.81, 0.78, and 0.76 for ATK inhibitor VIII.1, PIK-93, and GSK690693, respectively. Similar results were observed for other molecularly targeted drugs, such as anti-EGFR drugs (**Supplementary Fig. S3**). The results indicate that ResGitDR learns to predict drug response in a cell-state-oriented manner instead of relying on the genomic status of the biomarker genes. **Table. 1** shows the important SGAs gene in top 50 features in different pathways when predicting the drug response. For instance, in the PI3K/MTOR pathway, PIK3CA and PTEN are identified as important genes. On the other hand, in the ERK MAPK pathway, BRAF is recognized as a significant gene.

**Table 1.**
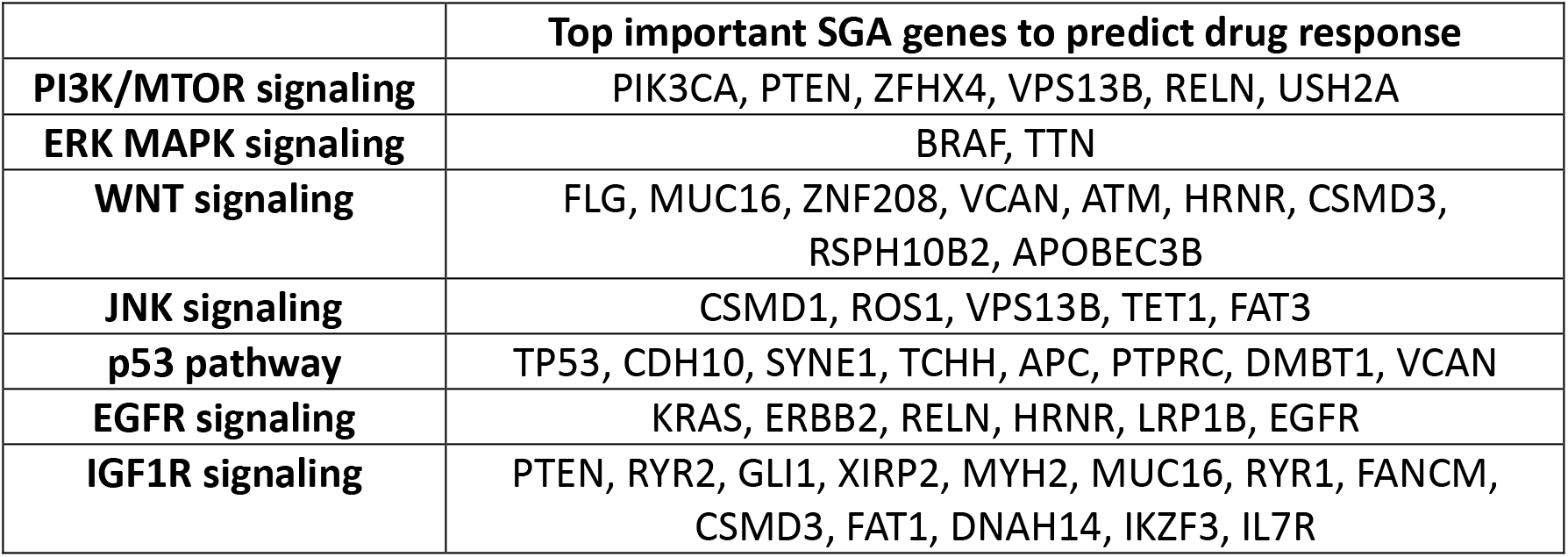
The top important SGA genes that were included as features when predicting the response to specific signaling pathways drugs.

We further investigated the cell-state-oriented nature of ResGitDR from another perspective. If a family of drugs shares a common MOA, it is expected that they will have a similar impact on cells sharing similar cell states. For each drug, we extracted the parameter vectors of the elastic net in ResGitDR model, which reflect the relative importance of features used by the model. We call this representation "drug embeddings", and we performed pairwise cosine similarity analysis of the drug embeddings. For each drug, we identified five drugs with the closest embeddings and visualized the relationships among the drugs (**Fig. 5D**). The results show that drugs targeting a common pathway share similar embeddings, supporting our assumption that ResGitDR identified the features reflecting the cell states indicative of sensitivity to drugs sharing MOAs.

## Discussion

In this study, we presented a novel framework for genome-informed precision oncology. Our approach overcomes the limitations of the current rule-based precision oncology ^5,8^ or simple machine learning approaches of directly using SGAs as inputs to predict drug responses ^9^. Instead, we designed the biologically-motivated ResGit model that learns to encode the information of SGAs with respect to gene expression using hierarchically organized latent variables, which mimic the cellular signaling systems of cancer cells. Hence, by transforming genomic data into features reflecting the functional state of cellular signaling systems, the integrated ResGitDR achieved significantly enhanced performance in predicting drug response.

Several novel designs in ResGitDR contribute to its utility. First, ResGit closely mimics the processes by which SGAs perturb cellular signaling systems, eventually leading to cancer. The cellular signaling system consists of hierarchically organized signaling proteins, and genomic perturbation at the different levels of the hierarchy exert distinct effects on cellular systems. By connecting SGAs to all latent variables, ResGit can learn the direct impact of an SGA on the specific components of the signaling system and allow the neural network to transmit such impact through the system. This makes the system transparent and interpretable, enabling ResGit to capture more efficiently the shared functional implications of different SGAs that perturb a common pathway in cells. Second, the self-attention mechanism enables the ResGit to capture the instance-specific impact of SGAs on the cellular signaling system, enabling the model to detect different roles of SGAs in individual tumors. Finally, explicitly including the hidden representation in ResGitDR makes the state of latent variables transparent, which enables ResGitDR to perform drug response prediction in a cell-state-oriented fashion.

Conventional genome-informed precision oncology mainly uses genomic biomarkers to guide the application of molecularly targeted drugs. As pointed out in previous studies ^5,9^ and our experiments, the accuracy of the rule-based or simple “black box” neural net models for guiding molecularly targeted drugs has room to be improved. Here, we show that by learning a representation of the cell signaling system, ResGitDR significantly outperforms simple models such as elastic networks and feed-forward neural networks. Although the current model has limited clinical utility because it is trained with pre-clinical data and not tested in real-world patient data, we anticipate that our framework has the potential to improve the accuracy of genome-informed targeted therapy in clinical settings if trained with large real-world data. Moreover, our framework can be expanded to guide chemotherapies as demonstrated by our results and other studies ^24-26^, which will significantly expand the scope of precision oncology beyond the genome-informed application of molecularly targeted drugs.

## Materials and methods

### Somatic genomic alterations (SGAs) pre-processing

The mutation data of GDSC was downloaded from Iorio *et al.* ^9^ and the CNV data and cancer type data were downloaded from Cell Model Passports (https://cellmodelpassports.sanger.ac.uk). The mutation data of TCGA were downloaded from the TCGA website (https://portal.gdc.cancer.gov), and the CNV data and cancer type data were downloaded from the Xena portal (http://xena.ucsc.edu). We represent an SGA event in a gene in a tumor as a binary variable, such that genes with mutations or somatic copy number alteration (deletion or amplification) were given a value of 1 and otherwise were given a value of 0. Since the majority of SGAs observed in tumors are likely passenger events, we take the union of 527 driver genes defined by the Cell Model Passports, 634 genes that are found to causally influence gene expression in cancers identified by Cai et al ^27^, and 324 mutation genes used in Foundation Medicine (https://www.foundationmedicineasia.com) to obtain the final set of 1,084 SGAs.

### Gene expression and TF-target gene matrix pre-processing

To take advantage of existing cancer big data, we combined both TCGA and GDSC RNA-Seq data. The RNAseq data of GDSC was obtained from Garcia-Alonso *et al.* ^28^ and of TCGA from the Xena portal. We selected the genes using the gene set described in Ding *et al.^24^* with the selection rule that genes with high variances were identified by medium variance analysis, bimodal mixture fitting, and statistical significance of modes. We obtained the processed TF-gene connectivity matrix from Tao *et al* ^15^. If a TF is known to regulate a gene, the corresponding element in the connectivity matrix is 1; otherwise, it is 0. The final set contained 320 TFs and 1,613 genes and had 105,224 connections.

### Drug sensitivity data pre-processing

Drug sensitivity data were downloaded from the GDSC website (https://www.cancerrxgene.org), and activity area (AA) was used to evaluate drug responses. In the GDSC1 dataset, there are a total of 367 drugs. Within this dataset, there are multiple drugs that share the same name but have different drug IDs. We considered these drugs as distinct entities. To facilitate future application in clinical practice, we discretized the drug response of a cell line with respect to a drug into two categories, sensitive (1) and resistant (0), by applying the waterfall method to each drug which was described in Ding *et al.^24^.* Specifically, the drug sensitivity measurements of all cell lines to a specific drug are sorted to generate a waterfall distribution. A linear regression is fitted to this distribution, and a Pearson correlation determines the goodness of fit. If the correlation coefficient is <0.95, the major inflection point is estimated as the point with maximal distance from a line drawn between the start and end points. If the correlation coefficient is >0.95, the median value is used. This value serves as the cutoff to separate sensitive and resistant cell lines to this drug.

### ResGitDR architecture

The overall architecture of ResGitDR is shown in **Fig. 1**. The model has two modules: 1) The Representation Learning Module (ResGit), which is a deep learning model that aims to encode the impact of SGAs on cellular signaling system by performing the task of predicting gene expression using SGAs and cancer type data as input. When trained, the model can be used to infer the state of the cellular signaling system by feeding SGAs and cancer type into the model. 2) The Drug Response Prediction Module, which utilizes elastic net to predict drug sensitivity by taking the hidden features learned in the first module and SGAs as input.

### Representation Learning Module in ResGitDR

The residual genomic impact transformer (ResGit) is similar to the genomic impact transformer (GIT) model developed by Tao *et al.*^29^ with several modifications of the architecture and procedures. Compared with GIT model, ResGit has more than one hidden layer and allows the connection of the SGAs to both the first hidden layer and each additional hidden layer (**Fig. 1C**). Through a series of hyperparameter tuning experiments, we set the number of hidden layers in ResGit to 4 (*H*_1_, *H*_2,_ *H*_3,_ and *TF*) and the number of hidden nodes number in *H*_1_, *H*_2,_ *H*_3,_ and *TF* layers to 200, 200, 200, and 320, respectively.

Input to the model consists of the cancer type label and *m* SGAs observed in a tumor. The inputs is firstly converted into embeddings using the "torch.nn.Embedding" class in PyTorch. The cancer type of the sample is transformed into a cancer-type embedding (*e*_*c*_) through an embedding layer. To capture the diverse impacts of a specific gene *m* on different hidden nodes, four distinct embedding layers are employed to convert the SGA gene *m* into four embedding vectors 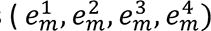. Additionally, instead of randomly initializing the SGA embeddings, we applied the Word2Vec^30^ algorithm to the SGA data to “pre-train” the SGA embedding. Embeddings learned in this fashion can capture the co-occurrence patterns of SGAs, so that the SGAs affecting a common pathway share a similar embedding. After initializing the SGA embedding with the pre-training gene embedding, the SGA embedding will further update with the supervision of gene expression data in ResGit.

After obtaining SGAs embedding, we employed a multi-head self-attention mechanism, which could distribute importance weights to SGAs in the training phase. Given a specific sample with cancer type (*C*) and a set of SGAs events (*M*), we obtained the first signal embedding layer (*e_1_) by the Equation (1)*, then applied a Relu activation function to get the first hidden representation (*H_1_*) through Equation (2):

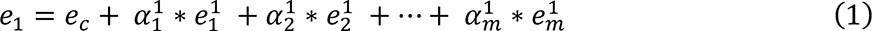

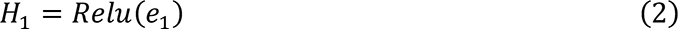

Where 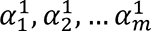 are the attention weights for the first hidden layer.

The attention weights in our experiment were calculated using the method described in Tao *et al^29^.* In brief, we calculated the attention weights 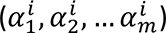 for hidden layer *H_i_* by following steps. First, the single-head (*h*) attention weights were calculated by Equation (3):

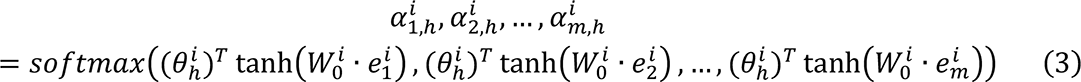

Where 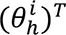 is the single-head parameter for head *h* and 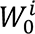 is the parameter matrix, both of them are for hidden layer *H_i_*. Then we calculated the multi-head attention weights by adding all the single head weights:

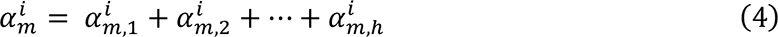

To obtain the subsequent signal embedding layer (*e*_2_-*e*_4_), only SGAs were used:

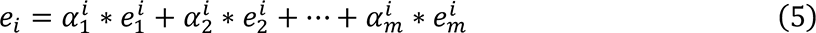

In order to obtain the second hidden representation layer (*H_2_*), we performed an addition operation to combine the initial hidden representation layer (*H_1_*) with the signal embedding layer (*e*_2_). Subsequently, we applied a ReLu layer:

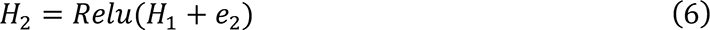

Similarly, we obtained the third hidden representation layer (*H*_3_):

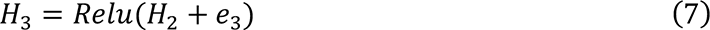

To obtain the last hidden layer, the transcription factor layer (TF layer), we used sigmoid function instead:

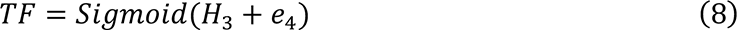

We used the TF layer to represent the state of transcription factors (TFs) explicitly, and the last linear layer learns the relationships between TFs and their target genes. This learning process was guided by a sparse matrix of prior knowledge derived from a TF-gene connectivity matrix (*P*∈ ^24k×l^), where *k* is number of TF and *l* is the number of gene. To predict the gene expression values, the Equation (9) was used:

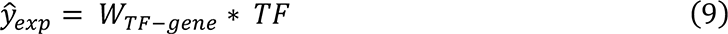

Where 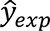 is the predicted gene expression value, *W*_*TF*–*gene*_ share the same shape with prior matrix P, and *W*_*TF*–*gene,i,j*_ is allowed to be nonzero and updated during learning only when P_i,j_ = 1. The gene expression is a continuous value, and mean square loss was used as the loss function:

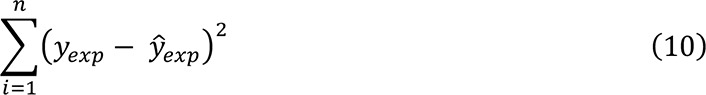

Where *n* is the number of samples, and *y*_*exp*_ is the observed gene expression value.

To avoid overfitting and increase robustness, we applied the pruning technique on both hidden layers and gene embeddings. For the weights matrix of the first three hidden layers, 90% of low-ranking weights are removed. For the last TF-gene expression weights matrix, we used prior knowledge, the TF-to-gene matrix, to regulate the weights, and the connections in this matrix are about 20%. For the embedding pruning, for each layer, every gene has its own gene embedding (the dimension of the first three layers is 200, and of the last one is 320). We first train the ResGit model without pruning any element in embedding, and after the model converges, we rank the nodes of each embedding, only nodes with the top 60% high value will be kept, and other elements will be changed into zero. Then, we re-train the ResGit module again till it converges. We used 10-fold cross-validation to evaluate the performance.

### Drug Response Prediction Module in ResGitDR

We used the elastic network model as the classifier for ResGitDR, which is a form of logistic regression with a hybrid regularization term that combines lasso and ridge regularization. We concatenated the original SGAs and latent variables derived by ResGit (*H*_1_, *H*_2_, *H*_3_, *TF*) of cell lines as the input features for the classifier, and binary drug-sensitivity label as targets. We used class sklearn.linear_model.LogisticRegression with penalty of elastic net. It contains two hyperparameters, L1_ratio and *C*. L1_ratio defines the relative weight of the lasso and ridge penalization terms, and *C* determines the regularization strength. We used grid search to select L1_ratio and *C* for each drug. The elastic net was performed with 10-fold cross-validation. Since ResGitDR model contains two modules, to avoid data leakage, we performed the cross-validation experiment simultaneously, using the same training/testing dataset for ResGit and elastic net.

To predict drug sensitivity, *SGAs*, *H*_1_, *H*_2_, *H*_3_, *TF* were firstly concatenated together, then elastic net was used:

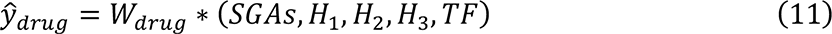

Where 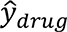 is the predicted drug sensitivity value and *W_drug_* is the weight matrix of elastic net.

Since drug response is binarized, the cross-entropy loss was used as loss function:

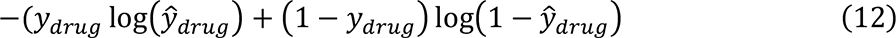

Where *y_drug_* is the observed drug sensitivity value.

## Abbreviations

ACC: Adrenocortical carcinoma
ALL: Acute lymphoblastic leukemia
BLCA: Bladder Urothelial Carcinoma
LGG: Brain Lower Grade Glioma
BRCA: Breast invasive carcinoma
CESC: Cervical squamous cell carcinoma and endocervical adenocarcinoma
CHOL: Cholangiocarcinoma
CLL: Chronic Lymphocytic Leukemia
COAD/READ: Colon adenocarcinoma/Rectum adenocarcinoma
DLBC: Lymphoid Neoplasm Diffuse Large B-cell Lymphoma
ESCA: Esophageal carcinoma
GBM: Glioblastoma multiforme
HNSC: Head and Neck squamous cell carcinoma
KICH: Kidney Chromophobe
KIRC: Kidney renal clear cell carcinoma
KIRP: Kidney renal papillary cell carcinoma
LAML: Acute Myeloid Leukemia
LCML: Chronic Myelogenous Leukemia
LGG: Brain Lower Grade Glioma
LIHC: Liver hepatocellular carcinoma
LUAD: Lung adenocarcinoma
LUSC: Lung squamous cell carcinoma
MB: Medulloblastoma
MESO: Mesothelioma
MM: Multiple Myeloma
NB: Neuroblastoma
OV: Ovarian serous cystadenocarcinoma
PAAD: Pancreatic adenocarcinoma
PCPG: Pheochromocytoma and Paraganglioma
PRAD: Prostate adenocarcinoma
SARC: Sarcoma
SCLC: Small Cell Lung Cancer
SKCM: Skin Cutaneous Melanoma
STAD: Stomach adenocarcinoma
TGCT: Testicular Germ Cell Tumors
THYM: Thymoma
THCA: Thyroid carcinoma
UCS: Uterine Carcinosarcoma
UCEC: Uterine Corpus Endometrial Carcinoma
UVM: Uveal Melanoma

## Acknowledgments

This study is supported by the NIH grant 5R01LM01201.

## Supplementary Figures

**Supplementary Fig. S1.**
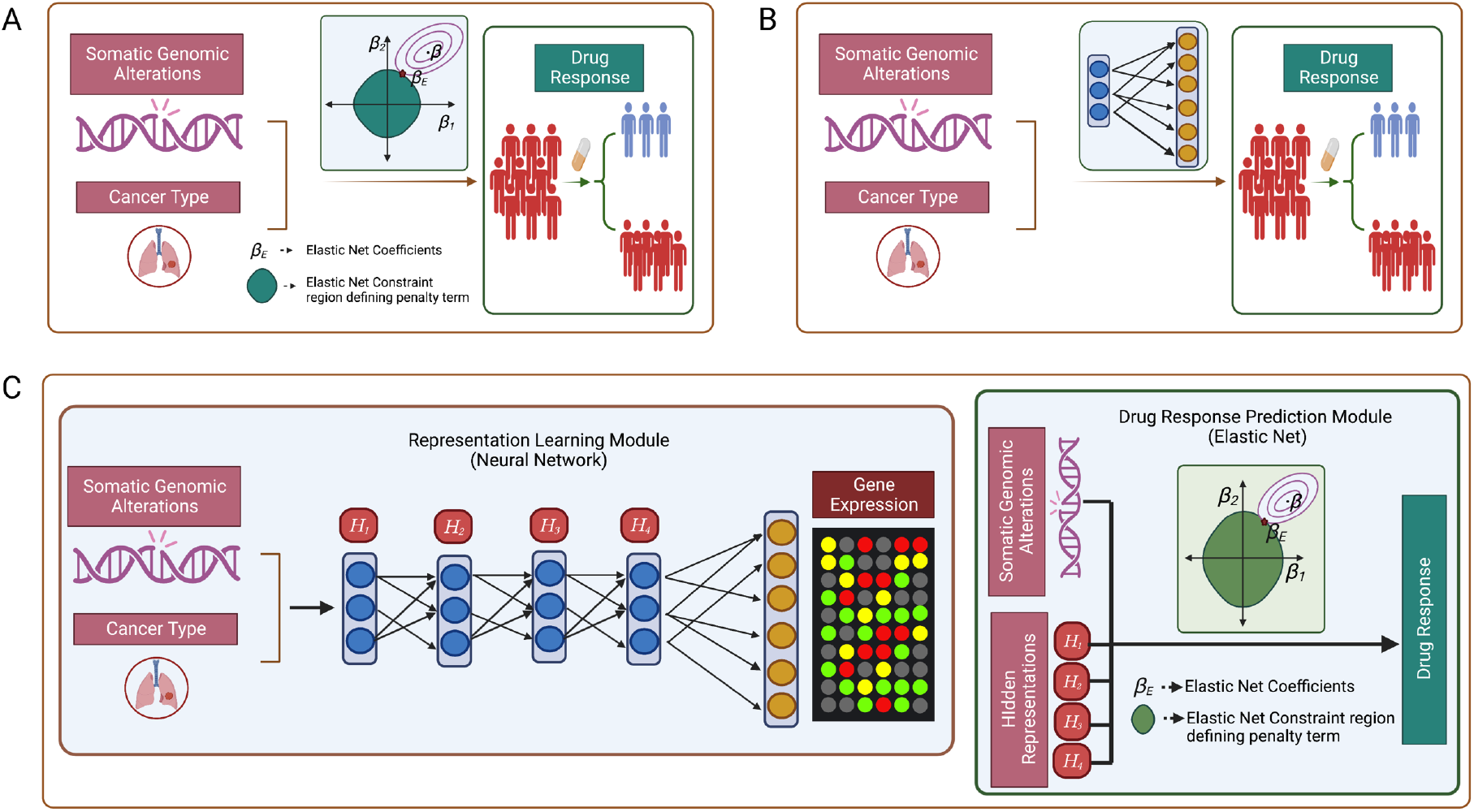
The model architectures of **(A)** the elastic net (EN) and **(B)** neural network (NN) models. Both models take SGAs and cancer type as inputs to directly predict drug response. **(C).** The architecture of the NNDR model involves a four-layer neural network (NN) that predicts gene expression using cancer type and SGAs as input. In the drug prediction phase, the NN is used to infer the state of hidden nodes, which are further used as inputs for the drug response prediction model.

**Supplementary Fig. S2.**
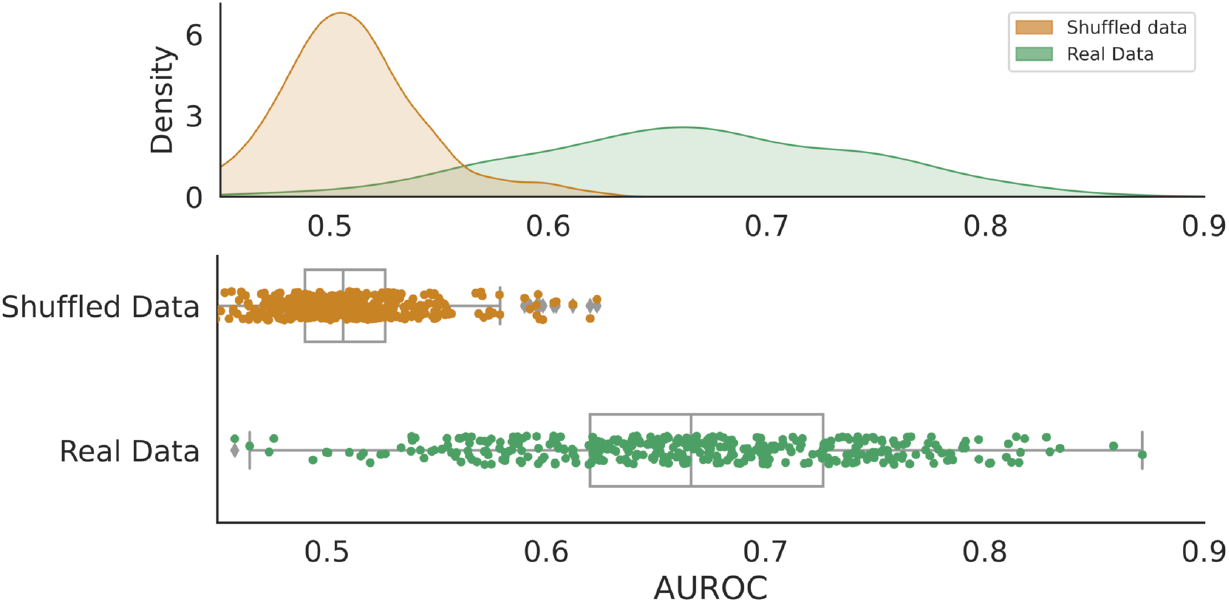
Using ResGitDR to predict drug sensitivity with shuffled data and Real Data

**Supplementary Fig. S3.**
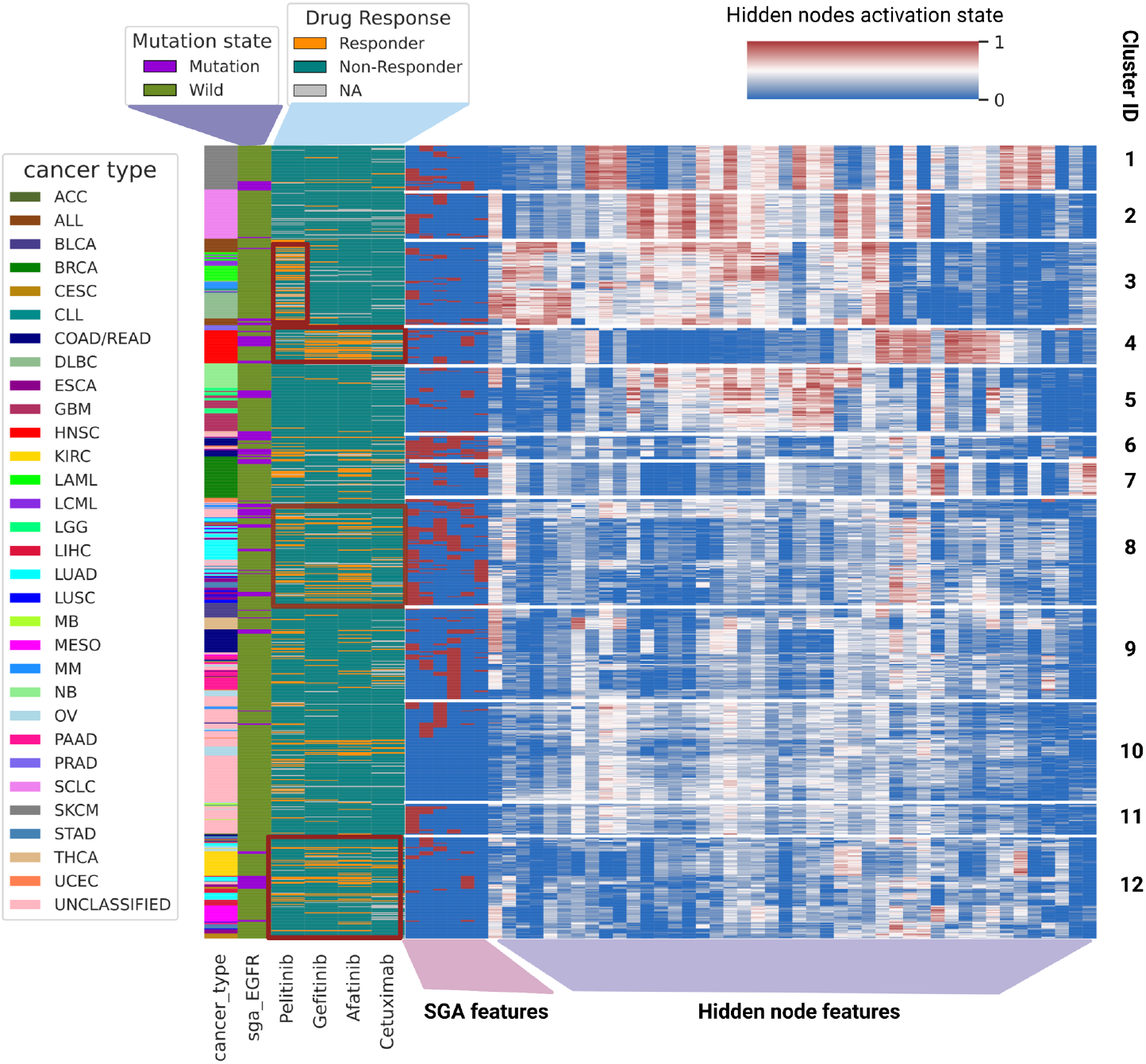
Cell-state-oriented prediction of sensitivity to anti-EGFR drugs. Annotations are the same as Fig. 5 in the main text.

## Supplementary Table

**Supplementary Table S1.** The targeted therapy drugs and chemotherapy drugs with AUROC higher than 0.7 when using ResGitDR, NNDR and EN to predict drug response (please see the excel file).

